# Drug targeting Nsp1-ribosomal complex shows antiviral activity against SARS-CoV-2

**DOI:** 10.1101/2021.11.02.466951

**Authors:** Mohammad Afsar, Rohan Narayan, Md Noor Akhtar, Huma Rahil, Sandeep M Eswarappa, Shashank Tripathi, Tanweer Hussain

## Abstract

The SARS-Cov-2 non-structural protein 1 (Nsp1) contains an N-terminal domain and C-terminal helices connected by a short linker region. The C-terminal helices of Nsp1 (Nsp1-C-ter) from SARS-Cov-2 bind in the mRNA entry channel of the 40S ribosomal subunit and block the entry of mRNAs thereby shutting down host protein synthesis. Nsp1 suppresses the host immune function and is vital for viral replication. Hence, Nsp1 appears to be an attractive target for therapeutics. In this study, we have *in silico* screened Food and Drug Administration (FDA)-approved drugs against Nsp1-C-ter and find that montelukast sodium hydrate binds to Nsp1-C-ter with a binding affinity (K_D_) of 10.8±0.2 μM *in vitro* and forms a stable complex with it in simulation runs with a binding energy of −76.71±8.95 kJ/mol. The drug also rescues the inhibitory effect of Nsp1 in host protein synthesis as demonstrated by the expression of firefly luciferase reporter gene in cells. Importantly, montelukast sodium hydrate demonstrates antiviral activity against SARS-CoV-2 with reduced viral replication in HEK cells expressing ACE2 and Vero-E6 cells. We therefore propose montelukast sodium hydrate may help in combatting SARS-CoV-2 infection.

## INTRODUCTION

SARS-CoV-2, the causative agent of severe coronavirus disease-19 (COVID-19) pandemic, is an enveloped positive-strand RNA-containing virus and belongs to the beta coronavirus family (V’Kovski et al., 2021). The virus contains nearly 30kb RNA genome with 5’-cap and 3’-poly-A tail (Finkel et al., 2021; V’Kovski et al., 2021). Upon entry into the host cell, the SARS-CoV-2 genome encodes for fourteen open reading frames (ORFs). The ORF1a and ORF1ab encode for two polyproteins, which are later auto-proteolytically cleaved into sixteen proteins, namely Nsp1 - Nsp16.

The cryo-electron microscopy (cryo-EM) structures of ribosomes from Nsp1-transfected human HEK293T cells indicate the binding of Nsp1 with the 40S and 80S ribosomal subunits (Schubert et al., 2020; Thoms et al., 2020; Tidu et al., 2020; Vankadari et al., 2020) (Figure 1A). Nsp1 contains 180 amino acids with N-terminal (1-127 amino acids) and C-terminal (148-180 amino acids) structured regions connected by a loop region of about 20 amino acids (Schubert et al., 2020; Thoms et al., 2020) (Figure 1B). This C-terminal region of Nsp1 (Nsp1-C-ter) contains two helices that harbours a conserved positively charged motif (KH-X_5_-R/Y/Q-X_4_-R). The deposition of positive charge towards one edge of these helices enhances their ability to bind the helix h18 of 18S rRNA. The other side of C-terminal helices interact with ribosomal proteins uS3 and uS5 in mRNA entry tunnel of 40S (Schubert et al., 2020; Thoms et al., 2020) (Figure 1A, zoomed view). These interactions enable Nsp1-C-ter bind deep into the mRNA entry tunnel and prevent the binding of mRNAs thereby inhibiting host protein synthesis (Schubert et al., 2020; Thoms et al., 2020; Tidu et al., 2020). Thus, Nsp1 helps in hijacking the host translational machinery (Yuan et al., 2020) and the cells are unable to mount innate immune response to counter the viral infection (Narayanan et al., 2008). Mutating the positively charged residues K164 and H165 in Nsp1-C-ter to alanines leads to a decrease in binding of Nsp1 with the ribosome and failure to inhibit host protein synthesis (Schubert et al., 2020; Thoms et al., 2020; Tidu et al., 2020).

**Figure 1:**
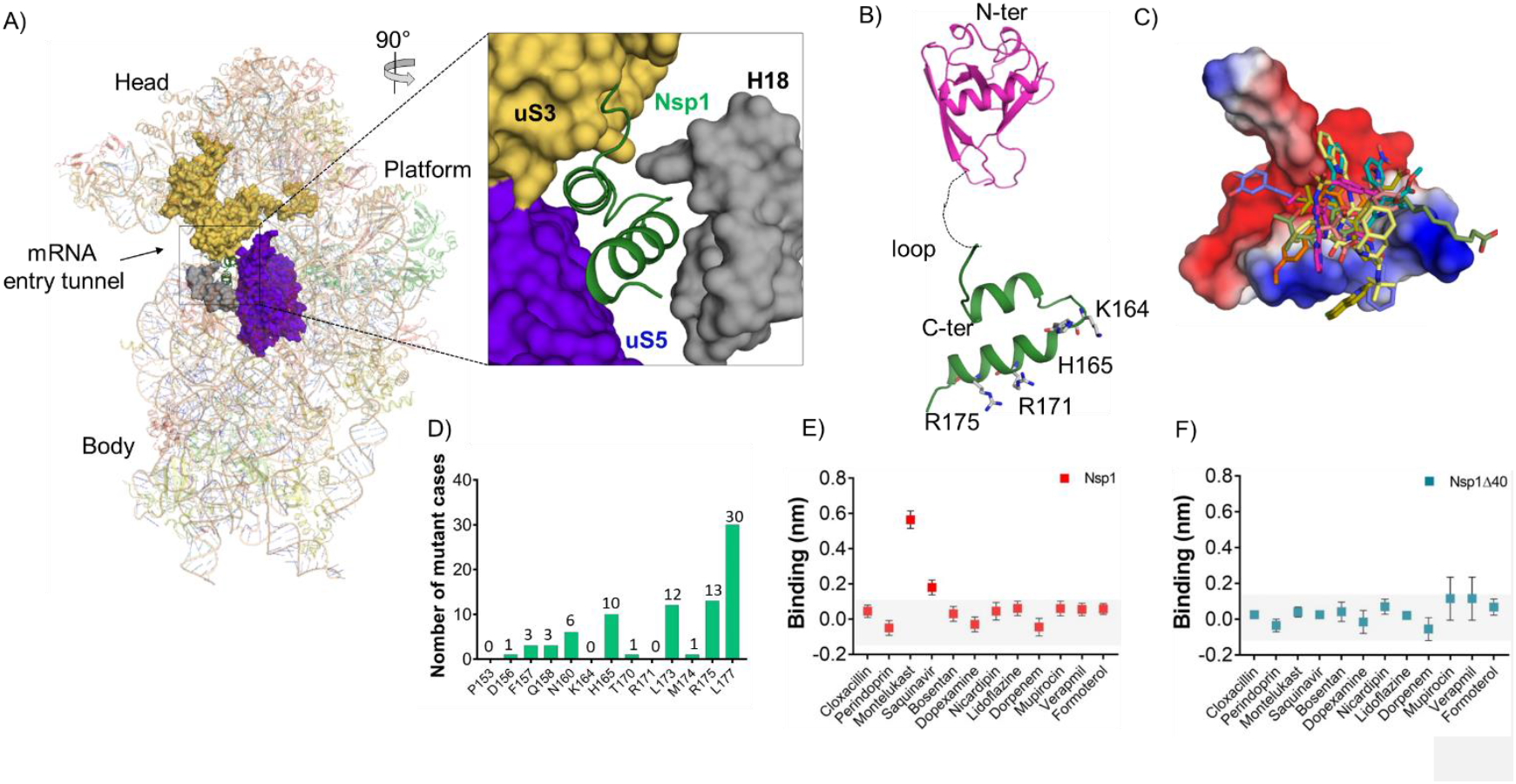
Screening of FDA-approved drugs against Nsp1 from SARS-CoV-2. A) The cryo-EM structure of the Nsp1-bound 40S ribosome (PDB:6ZOJ) shows the bound C-terminal helices of Nsp1 into the mRNA entry tunnel. The positively charged amino acids forms extensive interaction with h18 of 18S rRNA and the other side of the C-terminal helices interacts with uS3 and uS5. B) The structure of Nsp1 shows the presence of N-terminal structured region (PDB ID:7K7P) and C-terminal helices connected by a loop. C) Molecular screening of FDA-approved compounds led to identification of top hits. The docking mode of top hits (drugs) with Nsp1-C-ter is shown. D) The residues in Nsp1-C-ter involved in binding of selected drugs shows reduced mutational frequency. The analysis was performed on the worldwide deposited sequences of SARS-CoV-2 genome in GISAID database. The GISAID contains 4,440,705 genome sequences and we analyzed single nucleotide variants (SNV) for residues involved in drug binding. This analysis is performed with the help of GESS database (Fang et al., 2021). (E & F) BLI analysis for the initial screening of binding of the drugs with the (E) Nsp1 and (F) Nsp1Δ40 proteins.

Nsp1 is a highly conserved protein and less than 3% of the SARS-CoV-2 genomic sequences analysed showed mutation in Nsp1 (Min et al., 2020). Further, the Nsp1-C-ter showed a much reduced frequency of mutations (Min et al., 2020). The crucial role of the Nsp1 in inhibiting host gene expression, suppression of host immune response (Narayanan et al., 2008) and, notably, the reduced mutation frequency in Nsp1-C-ter across global SARS-CoV-2 genomes (Min et al., 2020) advocate targeting Nsp1 for therapeutics. In this study, we have employed computational, biophysical *in vitro* and *in vivo* studies to identify FDA-approved drugs targeting Nsp1-C-ter and check for its antiviral activity.

## Methods

### Receptor preparation for *in silico* studies and molecular screening of FDA-approved drugs

The three-dimensional coordinates of C-terminal helices of Nsp1 (Nsp1-C-ter; residue numbers 148 to 180) were taken from the cryo-EM structure of Nsp1-bound 40S (PDB ID: 6ZOJ). The close contacts, side chains, and bumps were fixed in Chimera (Pettersen et al., 2004). The molecule was minimized using the 100 steepest descent steps and ten conjugate gradient steps using the AMBERff14SB force field (Maier et al., 2015). None of the atoms were fixed during the minimization, and charges were assigned using the AMBERff14SB force field on standard residues. The final structure was optimized by the Powell method implemented in the biopolymer programme of SYBYL-X v2.1 (Tripos International, St. Louis, Missouri, 63144, USA).

The FDA-approved drug library was used to screen the drugs towards Nsp1-C-ter. The drug library containing 1645 compounds was subjected to molecular screening. The three-dimensional structure of (SDF format) compound library was optimized in the SYBYL-ligand prep module at default parameters. The single lowest strain energy tautomer for each compound was searched using the Surflex in the ligand preparation module. Subsequently, the binding pocket for ligands on Nsp1-C-ter was determined by the Computed Atlas of Surface Topography of proteins (CASTp) online server (Tian et al., 2018). The T151, P153, D156, F157, Q158, N160, K164, H165, S167, T170, R171, E172, L173, R175 and L177 were found to form the binding pocket. Finally, the library was screened against the 18S rRNA interacting interface of Nsp1-C-ter using the Surflex-dock program, which is available in SYBYL v2.1 (Jain, 2003). Twenty conformers were generated for each molecule with the 100 maximum rotatable bonds, and top potential molecules were selected based on docking score, which was calculated based on scoring function (flex C-score) using 48-core processor of HP Gen8 server.

### Nsp1 expression and purification

The gene construct encoding the Nsp1 from SARS-CoV-2 in pCDNA 5-3X-Flag-Nsp1 was amplified and sub-cloned into pET28a with N-terminal His-tag (Schubert et al., 2020; Thoms et al., 2020) using appropriate primers (Supplementary table 1). The sub-cloned construct was then further used to amplify and clone the C-terminal 40 amino acid deleted construct of Nsp1 (Nsp1Δ40) using appropriate primers (Supplementary table 1). Then the constructs were transformed into the *E. coli* BL-21 DE3 expression system. The secondary cultures were then inoculated with the 1% of the primary culture and incubated at 37°C at 180 rpm. After reaching the optical density (O.D.) of bacterial cells of 0.6 O.D., the cultures were then transferred at 16°C at 120 rpm, and the expression of pET28a-His-Nsp1 and pET28a-His-Nsp1Δ40 were induced by adding 1 mM of Isopropyl β-d-1-thiogalactopyranoside (IPTG) and allowed to grow for 18 hours. The bacterial cells were then harvested at 6000 rpm and resuspended in buffer A (50 mM HEPES-KOH pH 7.6, 500 mM KCl, 5 mM MgCl_2_, 5% Glycerol). The cells were then lysed by sonication at 18% amplitude and 10 sec on/off cycles for 10 min. The lysate was then subjected to centrifugation at 12000 rpm for 30 minutes to remove the cell debris. The clear supernatant was then loaded on the Ni-NTA beads (Qiagen) and incubated for 3 hours, and beads were then washed using buffer A. The bound protein was then eluted with buffer A supplemented with 300 mM imidazole, and purity was analysed on SDS-PAGE. The fractions which are containing the corresponding protein were concentrated and subjected to size exclusion chromatography on Superdex 200 increase 10/300 column in buffer B (50 mM HEPES-KOH pH 7.6, 150 mM KCl, 5 mM MgCl_2_, 2% Glycerol and 2 mM DTT). The pure protein fractions were pooled and concentrated between 2-8 mg/ml and stored in −80°C for further use.

### Drug-binding assays

#### Bio-layer Interferometry (BLI)

To identify the kinetic behaviour of the top selected compounds, we performed the label-free binding kinetics of the protein and ligands by using the bio-layer interferometry. The binding of ligand molecule on protein immobilized sensors induce change in the optical interference pattern of white light and is measured in nm. The Ni-NTA sensors were first activated by incubating in 10 mM Phosphate buffer saline for 10 min. After sensor activation, these sensors were then transferred to Buffer B. 2 μM of each protein was loaded on the Ni-NTA sensor and achieved a binding response of around 1 nm. The initial screening of the compounds was performed at 20 μM for all the *in silico* selected top hits. The drug molecules that showed binding response of more than 0.2 nm was selected for further kinetic experiments. Subsequently, the kinetic experiments were performed by incubating the protein-bound sensors with the increasing ligand concentration (0-25 μM) and binding was monitored. The data for control sensors (without protein) for each ligand concentration were also collected and subtracted from the response of proteins-bound sensors. The subtracted data was then analysed by fitting the 1:1 stoichiometric ratio for association and dissociation by applying the global fitting. Three independent experiments were performed to evaluate the steady-state kinetics and calculated K_D_ values.

#### Nanoscale Differential Scanning Fluorometry (NanoDSF)

*In silico* identified potential hits were then subjected to evaluate the binding with the His-Nsp1 and His-Nsp1Δ40 of SARS-CoV-2 protein. 2 μM of each protein was subjected to determine the melting temperature (Tm) in buffer B. The temperature scans were ranges from 20-90°C with the 1°C/min ramp size using Prometheus NT.48 NanoTemper. After successfully determining the melting temperature for individual proteins alone, we determined the ΔTm in the presence of drug molecules (10 μM) to identify the binding of the drug molecules. The top hits were selected for further evaluation in a change of the Tm by incubating with different concentrations of ligand (0-16 μM). The data was analysed by using ThermControl software.

### Molecular dynamics simulation of C-terminal helices of Nsp1 and drugs-bound complexes

The molecular dynamic simulations of FDA-approved drugs in complex with Nsp1-C-ter were selected based on the top binding score using BLI and NanoDSF. The final docked complexes were then prepared for molecular dynamics simulation studies. The systems for molecular dynamics studies were prepared for the Nsp1-C-ter alone and their complex with top hits using the Desmond v4.1implemented in Schrodinger-Maestro v11, where steric clashes and side-chain bumps were fixed. These prepared structures were then optimized by GROMOS96 54a7 force field (Schmid et al., 2011) and simple point charge water model was used to add the solvent molecules in the dodecahedron box with a distance of 1Å from the surface of the protein. Additionally, four sodium ions were added to neutralize the system. The following energy minimization was performed for all the systems with 5000 steps of steepest descent and conjugate gradient algorithms with threshold energy of 100 kCal/mol. The systems were then equilibrated in two phases, first is the isothermal-isochoric equilibration, where the constant number, volume, and temperature (NVT) was equilibrated for 100 picoseconds (ps), and the temperature of the system was monitored for all constants. In the second phase, the isothermal-isobaric equilibration was performed where the number of particles, pressure, and temperature (NPT) was equilibrated for 100 ps. After successful equilibration of the system, the final molecular dynamic runs were performed for 200 nanoseconds (ns) in three replicas with 2 femtoseconds of time steps. The root mean square deviation (RMSD), root mean square fluctuation (RMSF), and three-dimensional coordinates for all atoms of protein and ligands were extracted to analyse the molecular dynamics runs.

### Binding energy calculation

The binding energy for protein and ligands were calculated by applying the Molecular Mechanic and Poisson-Boltzmann Surface Area (MM-PBSA) (Genheden and Ryde, 2015; Wang et al., 2019). The two subsequent 100 ns runs from MD simulations were further subjected to perform the MM-PBSA by using the python script (mmpbsa.py) to calculate the binding energy of the two drugs.

This binding energy calculation quantitatively provides the *in silico* biomolecular interaction between the selected ligands and target protein. This binding energy mainly constitutes the polar solvation energy, non-polar solvation energy and potential energy. The free binding energy (ΔG_binding_) of the ligand was calculated by the following equation:

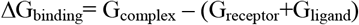

Where ΔG_complex_ describes the Gibbs free energy of the complex, G_receptor_ and G_lignad_ are total energy of protein and ligand, respectively.

### Luciferase-based assay: Translation inhibition and rescue experiments

We performed the luciferase based reporter assay to evaluate the target-specific action of the drug molecules. The HEK293 cells were transfected with 100 ng/well of pGL3-Fluc plasmid using Lipofectamine 2000 (Thermo Fisher Scientific) according to the manufacturer’s protocol at around 75-90% confluency in a 96 well plate. The plasmid expressing the Nsp1 protein (pcDNA 3.1-Nsp1) was co-transfected at 100 ng/well concentration. The transfection was performed in the presence of drugs montelukast and saquinavir at different concentrations. The cells were lysed 24 Hrs post-transfection, and luciferase activity was measured by using the Luciferase Reporter assay system (Promega Corporation) in the GLoMax Explorer system (Promega Corporation).

We further moved to check the expression level of *FLuc* and Glyceraldehyde 3-phosphate dehydrogenase (GAPDH). We isolated the total RNA from all conditions using the TRIzol by following the user manual protocol. Then we used 0.5 μg of total RNA to amplify the mRNA in the form of cDNA by using the RevertAid First Strand cDNA synthesis kit using manufacturer’s protocol. The amplified product was then used as template to amplify the FLuc and GAPDH gene in the presence of appropriate primers as mentioned in Table S1. The relative Ct values were monitored in the three replicates and relative fold change in expression was calculated. The significance of the data was monitored by applying the unpaired t-test through assuming Gaussian distribution parametric test by defining the statistical significance P<0.5.

To evaluate the total viral copy number, we isolated the RNA from SARS CoV-2 infected cells using TRIzol as per manufacturer‘s instructions, and equal amount of RNA used to determine the viral load using AgPath-ID™ One-Step RT-PCR kit (AM1005, Applied Biosystems). The primers and probes against SARS CoV-2 N-1 gene used are mentioned in Supplementary table S1. A standard curve was made using SARS CoV-2 genomic RNA standards, which was used to determine viral copy number from ct values.

### Cells and virus

The following cell lines were used in this study, namely HEK 293T cells (CRL-1573, ATCC, RRID: CVCL_0045), HEK 293T cells stably expressing human ACE2 (NR-52511, BEI Resources, NIAID, NIH, RRID: CVCL_A7UK), Vero-E6 cells (CRL-1586, ATCC, RRID: CVCL_0574). Cells were cultured in complete media prepared using Dulbecco’s modified Eagle medium (12100-038, Gibco) supplemented with 10% HI-FBS (16140-071, Gibco), 100 U/mL Penicillin-Streptomycin (15140122, Gibco) and GlutaMAX™ (35050-061, Gibco).

SARS-CoV2 (Isolate Hong Kong/VM20001061/2020, NR-52282, BEI Resources, NIAID, NIH) was propagated and quantified by plaque assay in Vero-E6 cells as described before (Case et al., 2020).

### Cytotoxicity assay

HEK-ACE2 cells were seeded in 0.1 mg/mL poly-L-lysine (P9155-5MG, Sigma-Aldrich) coated 96-well plate to reach 70-80% confluency after 24 Hrs. Vero-E6 cells were seeded in a regular 96 well plate to reach similar confluency. Cells were treated with 5, 10 and 20 μM montelukast or saquinavir in triplicates and incubated at 37°C/5% CO_2_. After 48 Hrs, cytotoxicity was measured using AlamarBlue™ Cell Viability Reagent (DAL 1025, Thermo Fisher) as per the manufacturer’s instructions.

### Western Blot

Cells were washed gently with 1X warm PBS (162528, MP Biomedicals), lysed using 1X Laemmli buffer (1610747, BIO-RAD), and heated at 95°C before loading on to a 10% SDS-PAGE gel. Separated proteins were transferred onto a PVDF membrane (IPVH00010, Immobilon-P; Merck) and incubated for 2hr with blocking buffer containing 5% Skimmed milk (70166, Sigma-Aldrich) in PBST (1X PBS containing 0.05% Tween 20 (P1379, Sigma-Aldrich)) for 2 hr at RT (room temperature). The blots were then probed with SARS-CoV-2 spike antibody (NR-52947, BEI Resources, NIAID, NIH) in blocking buffer for 12 hr at 4°C, followed by secondary Goat Anti-Rabbit IgG antibody (ab6721, Abcam, RRID:AB_955447) incubation for 2hr. Proteins were detected using Clarity Western ECL Substrate (1705061, BIO-RAD). Actin was labelled using antibody against beta-actin [AC-15] (HRP) (ab49900, Abcam, RRID: AB_867494). Relative intensity of bands was quantified using Fiji/imageJ.

### Virus infection

HEK ACE2 cells were seeded in poly-L-lysine coated 24-well plate to reach 80% confluency at the time of infection. Vero-E6 cells were seeded in a regular 24 well plate to reach similar confluency. Cells, in quadruplicates, were first pre-treated with 5 and 10 μM concentrations of montelukast sodium hydrate (PHR1603, Merck) or saquinavir mesylate (1609829, Merck) for 3hr in complete media, washed and infected with 0.1 MOI (HEK ACE2) or 0.001 MOI (Vero-E6 cells) SARS CoV-2. After 48 hr, cell culture supernatants were collected for plaque assay, and cells were harvested for western blot analysis or processed for total RNA extraction using TRIzol (15596018, Thermo Fisher). The drugs were present in the media for the entire duration of the experiment.

### Plaque Assay

Infectious virus particles from cell culture supernatants were quantified by plaque assay. Briefly, Vero-E6 cells were seeded in 12-well cell culture dishes, and once confluent, cells were washed with warm PBS and incubated with dilutions of cell culture supernatants in 100 μL complete DMEM for 1 hr at 37 °C / 5% CO2. The virus inoculum was then removed, and cells overlaid with 0.6% Avicel (RC-591, Dupont) in DMEM containing 2% HI-FBS. After 48 hr incubation, cells were fixed with 4% paraformaldehyde, and crystal violet (C6158, Merck) staining was done to visualize the plaques.

### Plasmids

pLVX-EF1alpha-SARS-CoV-2-nsp1-2xStrep-IRES-Puro expressing SARS CoV-2 NSP1 was a kind gift from Prof. Nevan Krogan (Gordon et al., 2020). Other plasmids used in this study include Plasmids pRL-TK (mammalian vector for weak constitutive expression of wild-type Renilla luciferase), pGL4 (mammalian vector expressing firefly luciferase), pIFN-β Luc (IFN beta promoter-driven firefly luciferase reporter). The plasmid pMTB242 pcDNA5 FRT-TO-3xFLAG-3C-Nsp1_SARS2 was a kind gift from Prof. Ronald Beckmann.

## RESULTS

Since repurposing a drug is a quicker way to identify an effective treatment, we screened FDA-approved drugs against Nsp1-C-ter (148-180 amino acids) which binds in the mRNA channel (Figure 1C). The drugs docked to a small region of Nsp1-C-ter consisting of residues (P153, F157, N160, K164, H165, and R171) which coincides with its ribosome-binding interface (Figure 1C). The residues in Nsp1-C-ter involved in binding drugs show minimal mutations in worldwide deposited 4,440,705 sequences of SARS-CoV-2 genome in GISAID database (Figure 1D). We identified top hits with at least three hydrogen bonds (H-bonds) near the ribosome binding site of Nsp1-C-ter (Table 1). Further, the clash that the drugs may have against the ribosome in its bound form with Nsp1-C-ter was also analyzed. Montelukast sodium hydrate (hereafter referred to as montelukast) and saquinavir mesylate (hereafter referred to as saquinavir) showed high clash scores (Table 1). Montelukast is regularly used to make breathing easier in asthma (Paggiaro and Bacci, 2011), while saquinavir is an anti-retroviral drug used in the treatment of human immunodeficiency virus (HIV)(Khan et al., 2021).

**Table 1.**
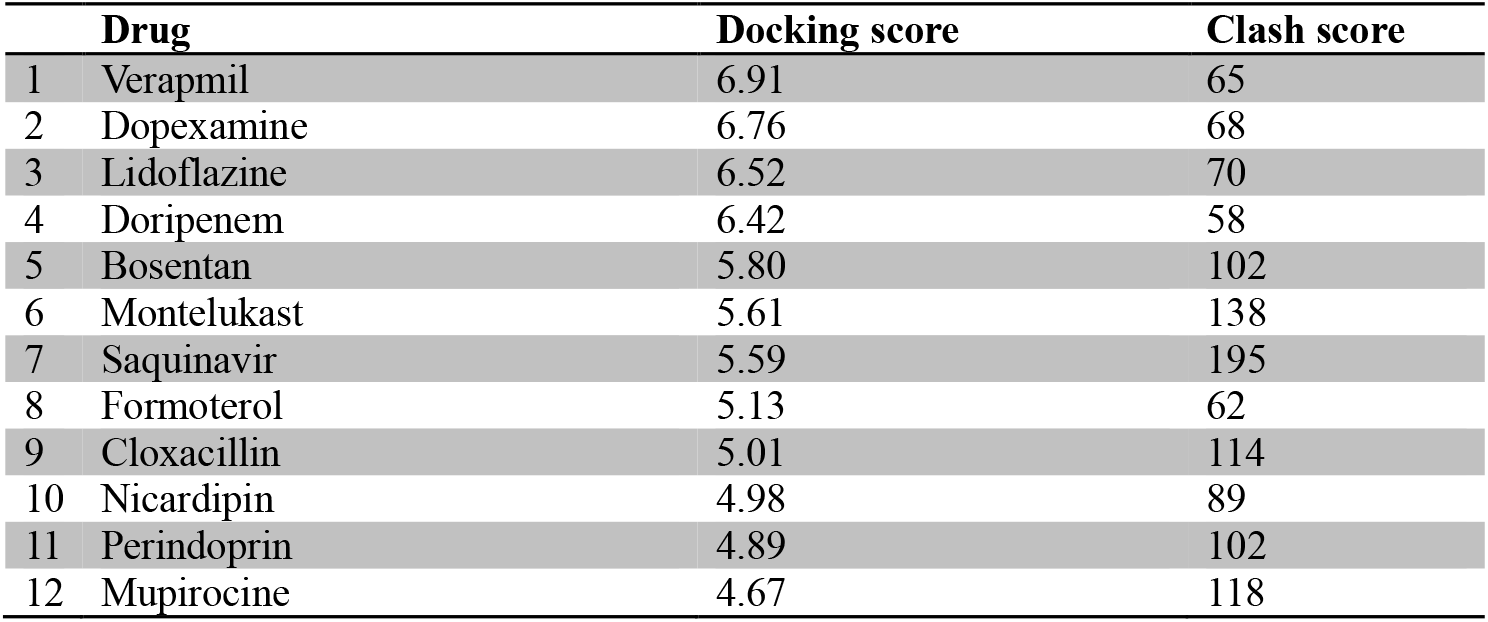
Top hits of FDA-approved drugs upon screening against Nsp1-C-ter.

|  | **Drug** | **Docking score** | **Clash score** |
| --- | --- | --- | --- |
| 1 | Verapamil | 6.91 | 65 |
| 2 | Dopexamine | 6.76 | 68 |
| 3 | Lidoflazine | 6.52 | 70 |
| 4 | Doripenem | 6.42 | 58 |
| 5 | Bosentan | 5.80 | 102 |
| 6 | Montelukast | 5.61 | 138 |
| 7 | Saquinavir | 5.59 | 195 |
| 8 | Formoterol | 5.13 | 62 |
| 9 | Cloxacillin | 5.01 | 114 |
| 10 | Nicardipin | 4.98 | 89 |
| 11 | Perindoprin | 4.89 | 102 |
| 12 | Mupirocine | 4.67 | 118 |

Next, all twelve drugs were tested *in vitro* for their ability to bind to Nsp1. The purified proteins, *i.e*., full-length Nsp1 and C-terminal helices truncated Nsp1 (Nsp1Δ40) proteins, were loaded on the Ni-NTA sensors in BLI, and the compounds were screened to determine its binding to these proteins. We found that montelukast and saquinavir show binding to Nsp1 (Figure 1E) but not with Nsp1Δ40 (Figure 1F). This indicates that montelukast and saquinavir bind to Nsp1-C-ter. The rest of the compounds does not show binding with Nsp1 or with Nsp1Δ40 (Figure 1E and 1F). We next determined binding affinities of montelukast and saquinavir against Nsp1. Montelukast shows a binding affinity (Kd) of 10.8±0.2μM (Figure 2A) while saquinavir shows a binding affinity of 7.5±0.5μM towards Nsp1-C-ter (Figure 2B).

**Figure 2:**
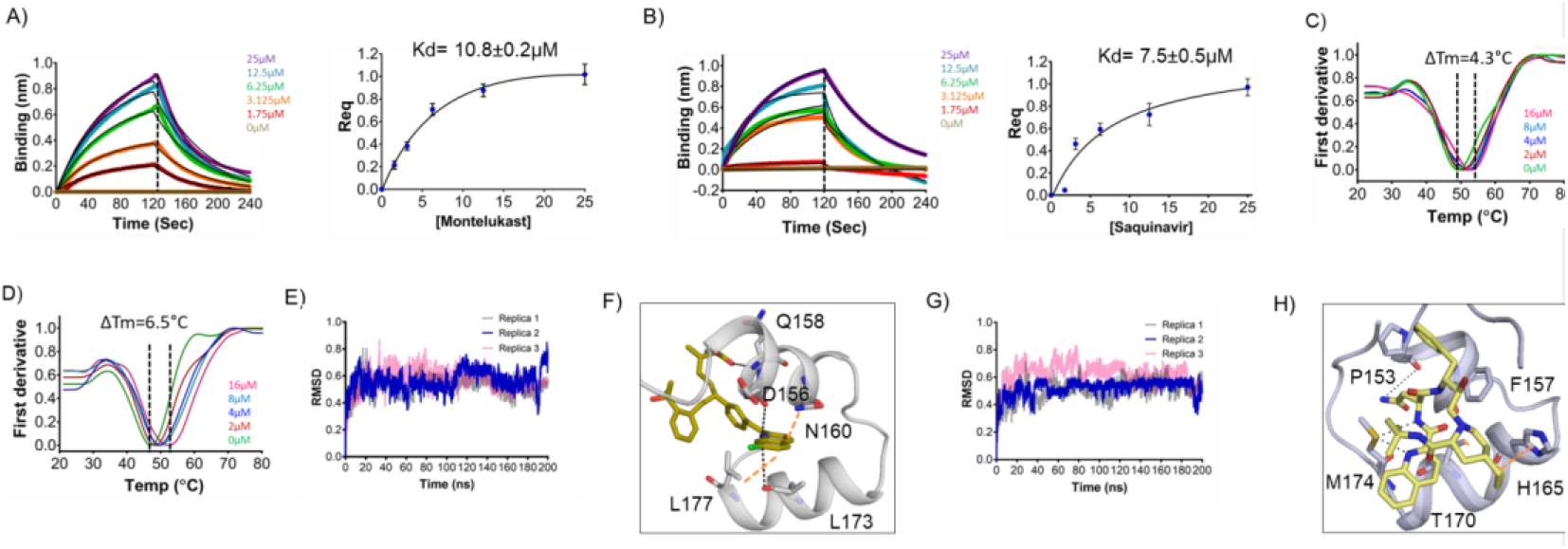
Binding kinetics and molecular simulation dynamics runs of drugs against Nsp1-C-ter. (A & B) The kinetic behaviors of (A) montelukast and (B) saquinavir monitored using BLI by incubating increasing concentration of the drug molecule (0-25μM) on the protein-bound sensors. Montelukast shows a binding constant (K_D_) of 10.8±0.8μM, while saquinavir binds with Nsp1-C-ter with a K_D_ value of 7.5±0.5μM. (Error bars represent standard deviation of three replicates in (A) and (B). (C & D) NanoDSF experiments to evaluate the change in the melting temperature of the Nsp1 by incubating increasing concentration of (C) montelukast and (D) saquinavir. (The experiments were performed in three replicates) E) Simulation runs with montelukast show stable RMSD values for all replica throughout all molecular dynamic simulation trajectories for 200ns. F) The analysis of binding mode of montelukast at the end of 200ns shows stable binding with C-terminal helices. The residues, D156, Q158 and L173, form H-bonds with montelukast, while N160 and L177 form base stacking interactions. G) Simulation runs with saquinavir show stable pattern in RMSD values throughout in all molecular dynamic simulation trajectories for 200ns. H) The analysis of binding mode of saquinavir at the end of 200ns shows stable binding with the C-terminal helices. The residues, P153 and M157, form H-bonds with saquinavir, while T170 and H165 form base stacking interactions.

To further validate the binding of ligands with Nsp1-C-ter, we performed NanoDSF experiments where we observed the change in the melting temperature of Nsp1 in the presence of drugs. We observed that only montelukast and saquinavir induce a change in the melting temperature of Nsp1 (Figure 2-figure supplement 1A). None of the ligands was able to change the melting temperature of the Nsp1Δ40 protein (Figure 2-figure supplement 1B). Next, we performed NanoDSF experiments with different concentrations of montelukast and saquinavir to determine the change in melting temperature of Nsp1. We observed that montelukast shifts the ΔT_m_ by 4.3°C while the saquinavir causes a ΔT_m_ shift by 6.5°C (Figure 2C and D). Overall, montelukast and saquinavir showed binding to Nsp1-C-ter *in vitro*.

To gain insights into the binding mode of montelukast and saquinavir with Nsp1-C-ter, we analyzed the docked drugs and performed molecular dynamic simulation runs. The molecular screening experiment shows the binding of montelukast with Nsp1-C-ter with a 5.61 docking score (Table 1 and Figure 2-figure supplement 2A). In the simulation runs the root mean square deviation (RMSD) of C-terminal helices bound with montelukast shows less deviation from the mean atomic positions (Figure 2E). The analyses of H-bonds and hydrophobic interactions indicate strong binding of montelukast during the simulation run. At the end of the simulation run, montelukast shows a stable complex by forming H-bonds with D156, Q158 and L173, while N160 and L177 form base stacking interactions (Figure 2F). The root mean square fluctuation (RMSF) plot shows the thermal stability of individual residues throughout the molecular dynamics run of the molecule, and it appears to be stable (Figure 2-figure supplement 2B). Saquinavir shows binding with Nsp1 with a docking score of 5.6 (Table 1 and Figure 2-figure supplement 2C). The RMSD plot of saquinavir bound C-terminal helices shows reduced deviation of the protein atoms during the simulation runs from the mean atomic position (Figure 2G). The residues P153, T170 and M174 form H-bonds with saquinavir while H165 forms base stacking interaction at the end of the run (Figure 2H). The RMSF plot show that the participating residues is also stabilised upon the binding of saquinavir (Figure 2-figure supplement 2D). Overall, the residues involved in binding montelukast and saquinavir show extremely low mutational frequency.

Furthermore, these drug-Nsp1 complexes were subjected to free binding energy calculations for 100 ns in two replicas for each complex. Montelukast and saquinavir bind with Nsp1 with binding energies of −76.71±8.95 kJ/mol and −72.46±3.34 kJ/mol, respectively (source data 3). The average H-bonds were analysed for the C-terminal region of Nsp1 alone and drug-bound complexes. We observed that these drugs-bound complexes show higher average H-bonds throughout different replica simulations (Figure 2-figure supplement 2E).

Since Nsp1 is known to inhibit host protein synthesis by blocking the mRNA entry tunnel on the ribosome and co-transfection of Nsp1 with capped luciferase reporter mRNA causes reduction of luciferase expression (Thoms et al., 2020). We hypothesized that binding of montelukast or saquinavir to Nsp1-C-ter may prevent inhibition of host protein synthesis. To test this hypothesis, we carried out the cell-based translational rescue of luciferase activity in the presence of montelukast and saquinavir in HEK293 cells when co-transfected with Nsp1. Co-transfection of Nsp1 decreased the luciferase activity by almost half, which is restored by the increasing amount of montelukast (Figure 3A). However, we do not observe a similar rescue of luciferase activity in the presence of saquinavir (Figure 3B). Further experiments are needed to figure out why saquinavir is unable to rescue the Nsp1-mediated translation inhibition. There was no significant change in gene expression of the firefly luciferase *FLuc* gene (Figure 3 C&D).

**Figure 3:**
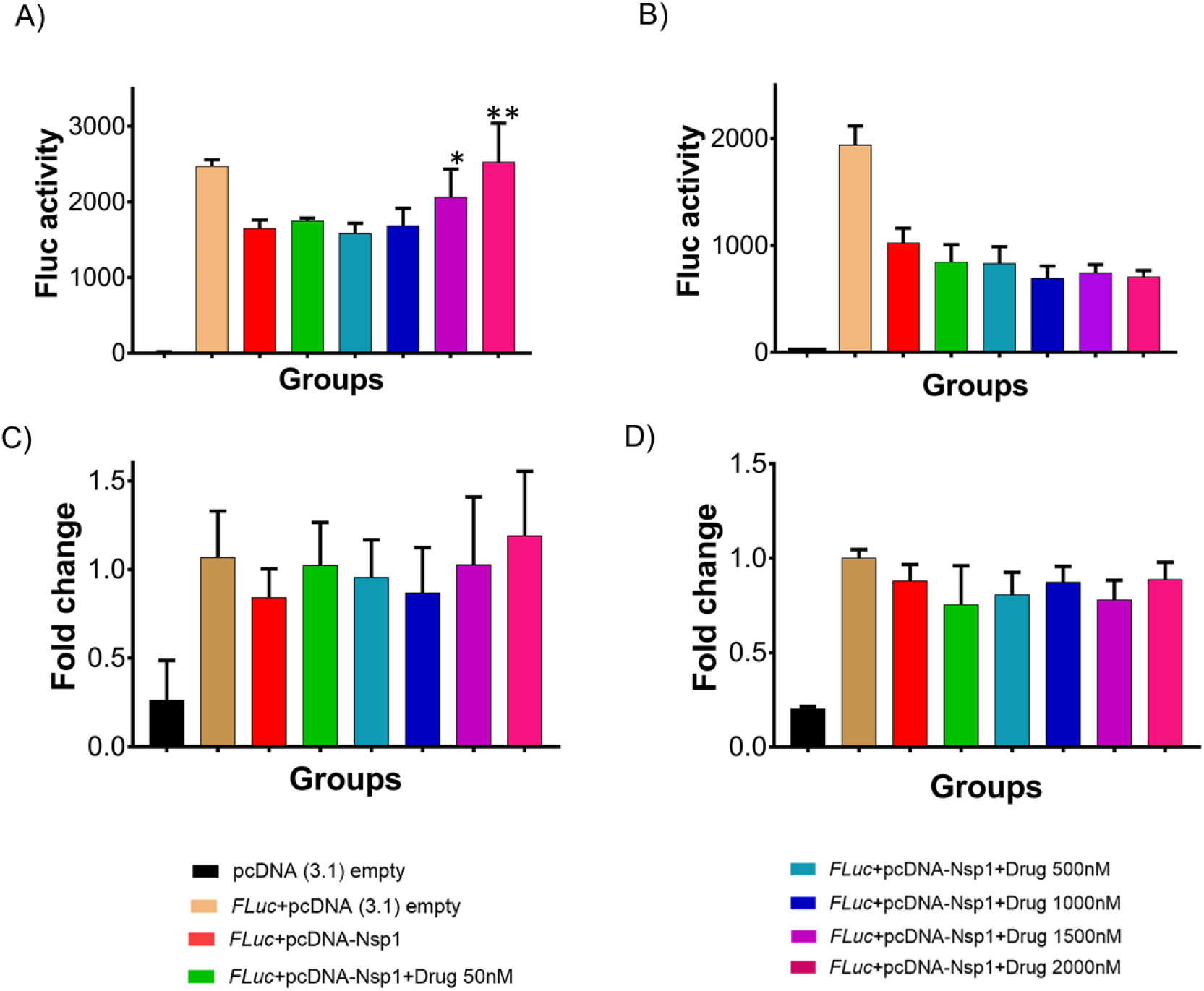
Translational rescue experiments in the presence of montelukast and saquinavir. A) Luciferase-based reporter assay shows translational rescue of luciferase in the presence of montelukast. B) Luciferase-based reporter assay shows that saquinavir could not rescue the luciferase expression. Error bars represent standard deviation of three replicates in (A) and (B). C & D) The real-time PCR to quantitate the fold change of *F Luc* gene in comparison to GAPDH in the presence of different concentration of the drug molecules. A) montelukast B) saquinavir. The panel below provides the details of experimental conditions. Error bars represent standard deviation of three replicates in (A) and (B). The significance of the data was monitored by applying the unpaired t-test through assuming Gaussian distribution parametric test by defining the statistical significance. **P < 0.01; ***P < 0.001; ****P < 0.0001. The error bars represent the standard deviation.

To test antiviral effects of montelukast and saquinavir against SARS CoV-2, we first tested the cytotoxicity of these drugs in HEK-ACE2 and Vero-E6 cells. Results showed minimal toxicity up to 10μM montelukast and saquinavir in both cell lines. However, in Vero-E6 cells, the highest concentration (20μM) of both drugs showed an almost 80% decrease in cell viability, compared to untreated cell control (Figure 4-figure supplement 1A&B). Based on this, a working concentration of 10μM or lower was used for both drugs. At a concentration of 10μM, montelukast showed significant antiviral activity, as indicated by reduced expression of viral spike protein in HEK-ACE2 and Vero-E6 cells (Figure 4A and D). The corresponding qRT-PCR data demonstrated up to 1-log reduction in viral copy number in both HEK-ACE2 and Vero-E6 cells at this concentration (Figure 4B and E), supported by a decrease in infectious virus titer measured by plaque assay (Figure 4C and F). No significant antiviral effects were observed in the presence of 10μM saquinavir (Figure 4-figure supplement 2 A-F).

**Figure 4:**
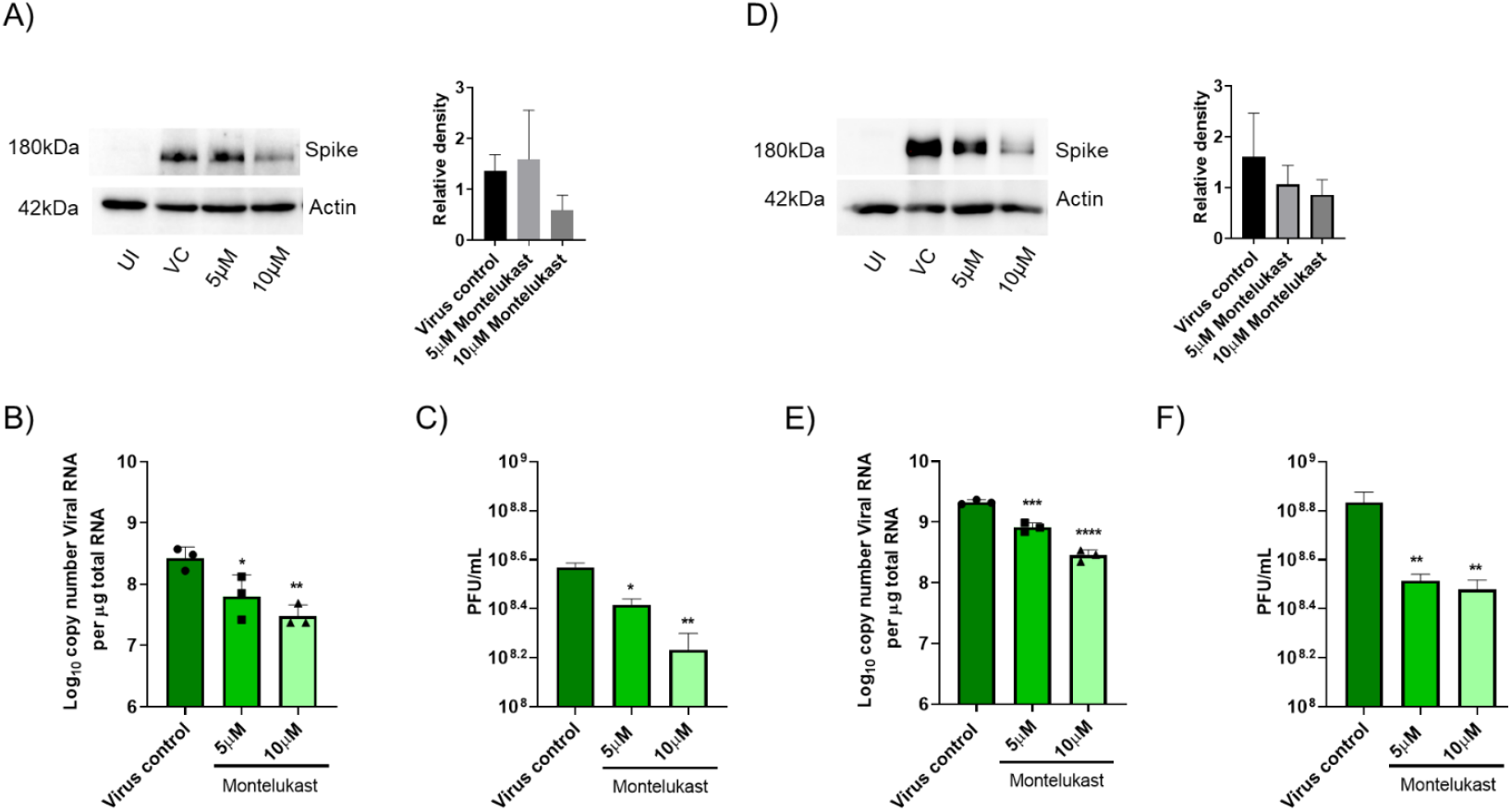
Montelukast shows antiviral activity against SARS-CoV-2. A) HEK ACE2 cells were pre-treated with 5 or 10μM montelukast and infected with 0.1 MOI SARS CoV-2 for 48hr. Virus spike protein expression by western blot analysis, with corresponding relative density of bands are shown. B and C) Viral RNA copy number from infected cells was quantified by qRT PCR, infectious virus titer from cell culture supernatants by plaque assay, respectively. Vero E6 cells were pre-treated with 5 or 10μM montelukast and infected with 0.001 MOI SARS CoV-2 for 48 hr. D) Virus spike protein expression by western blot analysis, with corresponding relative density of bands. E) Viral RNA copy number from infected cells was quantified by qRT PCR and F) infectious virus titer from cell culture supernatants by plaque assay. *P<0.05; **P<0.01; ***P<0.001; ****P<0.0001; ns-not significant, using one-way ANOVA with Dunnett’s multiple comparison test. Error bars represent standard deviation.

## DISCUSSION

Nsp1 binds to the 40S ribosomal subunit via its C-terminal helices in the mRNA entry tunnel, thereby blocking the entry of mRNAs leading to shutdown of host protein synthesis. Nsp1 mimics the binding mode of eIF3j, a non-stoichiometric subunit of eukaryotic initiation factor 3 (eIF3), which also binds in the mRNA entry tunnel and prevents the binding of the eIF3 complex (Babaylova et al., 2019; Cate, 2017; Sharifulin et al., 2016). However, Nsp1 does not inhibit the binding of cricket paralysis virus internal ribosome entry site (CrPV-IRES) mRNA to the ribosome, while it restricts the movement of the 40S head in the 48S pre-initiation complex (Yuan et al., 2020).

Since repurposing a drug is a quicker way to identify an effective treatment, we screened FDA-approved drugs against the Nsp1-C-ter and found montelukast as potential drugs against it. Montelukast is a leukotriene receptor antagonist and repurposing montelukast for tackling cytokine storms in COVID-19 patients has been suggested (Sanghai and Tranmer, 2020) and hospitalized COVID-19 patients that were given montelukast had significantly fewer events of clinical deterioration (Khan et al., 2021). Montelukast also appears as a hit against the SARS-CoV-2 main protease, (M^pro^) protease, in computational studies (Abu-Saleh et al., 2020; Sharma et al., 2021). However, Chunlong *et al*. demonstrated that montelukast gives false positive anti-protease activity as it cannot bind the GST-tagged-M^pro^ in thermal shift assay and native mass spectrometry experiments (Ma and Wang, 2021). Thus, montelukast may not be an inhibitor for M^pro^ protease.

Viruses employ different strategies to shutdown host translation machinery. In SARS-CoV-2, Nsp1 inhibits translation by binding to the mRNA channel. Here, we show that montelukast binds to Nsp1, rescues the Nsp1-mediated protein translation inhibition and has antiviral activity against SARS-CoV-2. The rescue of shutdown of host protein synthesis machinery by montelukast seems to contribute towards the antiviral activity of the drug; however, further experiments would be essential to figure out detailed mechanism of its antiviral activity. Overall, our study identifies C-terminal region of Nsp1 as a druggable target and montelukast as a potential antiviral drug against SARS-CoV-2 infection that may help in combatting the COVID-19 pandemic.

## Supporting Information

Supporting information contains 5 figures, 2 tables and details of materials and methods.

## Aknowledgements

This work was supported by Intermediate Fellowship from DBT-Welcome Trust India Alliance to TH (IA/I/17/2/503313). TH also thanks SERB for funds released under IRPHA (COVID-19 Life Sciences; File Number:IPA/2020/000094). ST acknowledges funding from DBT-BIRAC grant (BT/CS0007/CS/02/20) and DBT-Wellcome Trust India Alliance Intermediate Fellowship (IA/I/18/1/503613). We acknowledge Swarnajayanti Fellowship from DST to SME (SB/SJF/2020-21/18). The authors also acknowledge DBT-IISc Partnership Program Phase-II (BT/PR27952-INF/22/212/2018) for support.

## Notes

The authors declare no conflict of interest.

## Supplementary Information

**Figure 2 figure supplement 1:**
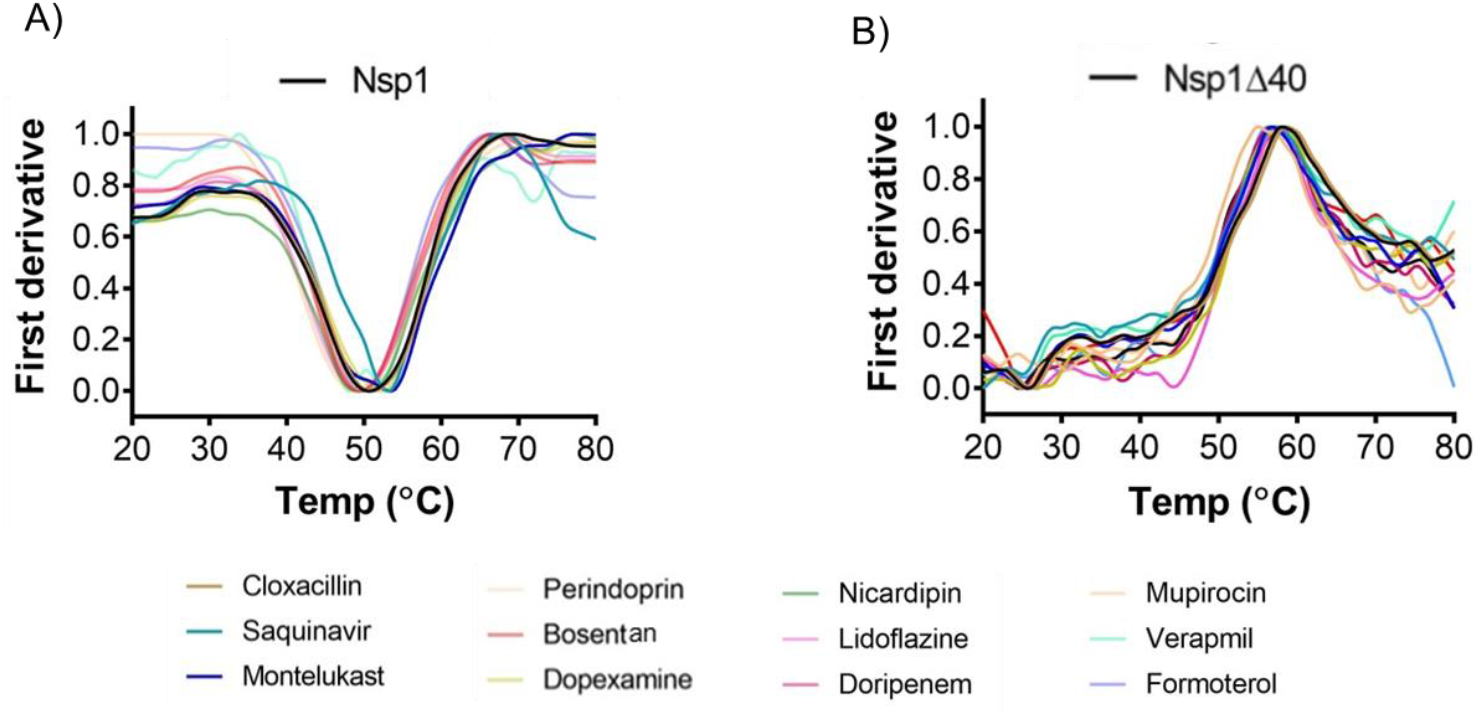
NanoDSF experiments to evaluate the binding of top hits with the Nsp1 and Nsp1Δ40. (A and B) The change in the melting temperature of (A) Nsp1 and (B) Nsp1Δ40 protein was monitored in the presence of the selected molecules. The melting curve for apo-proteins are shown in black color. Montelukast and saquinavir induce change in the melting temperature of Nsp1 while none of the molecules show any difference in the melting temperature of Nsp1Δ40 protein.

**Figure 2 figure supplement 2:**
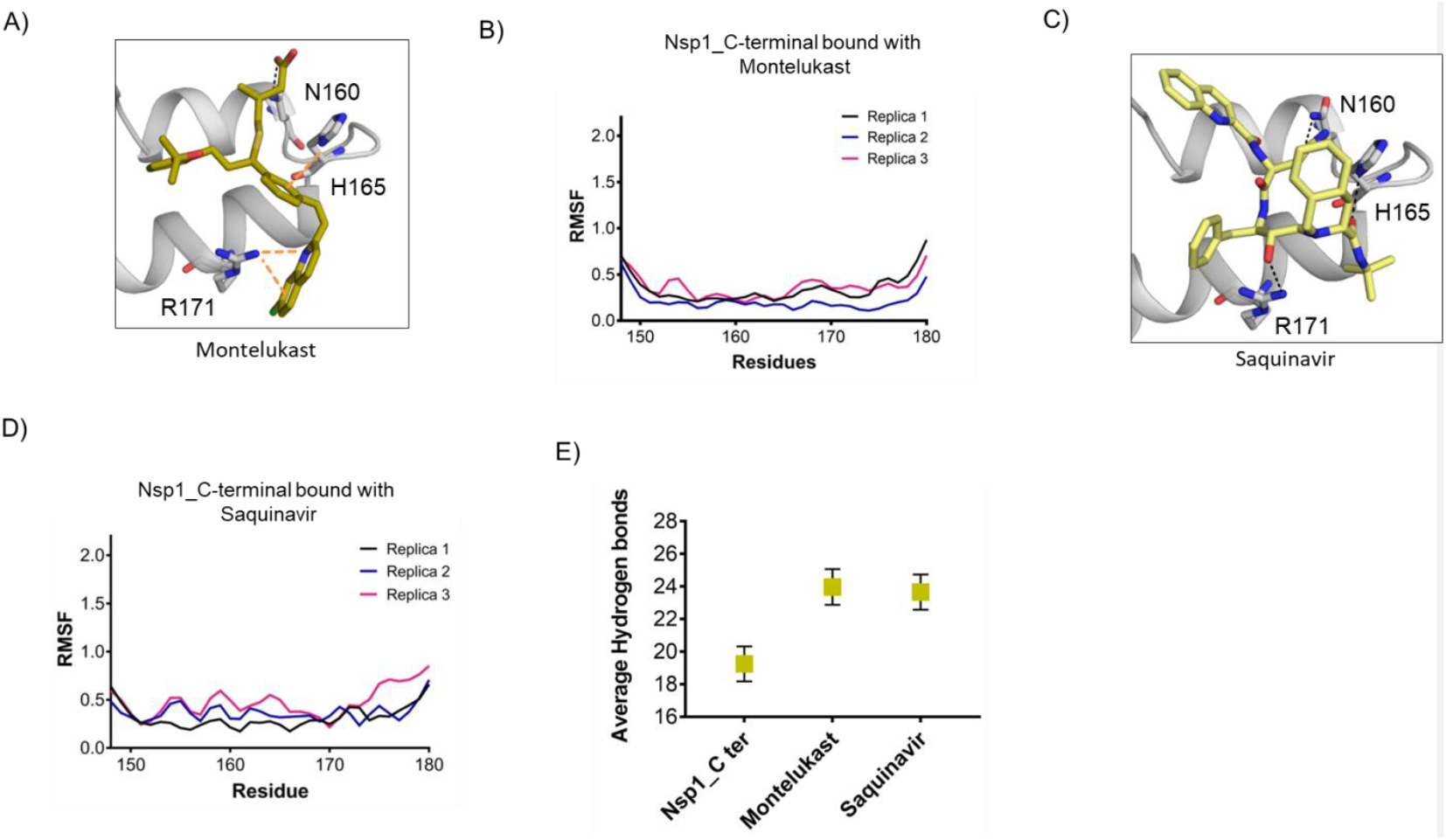
Structural dynamics of drug-bound complexes of Nsp1-C-ter. (A & B) The docking mode of (A) montelukast and (B) saquinavir with Nsp1-C-ter (C & D) The RMSF plot of (C) montelukast- and (D) saquinavir-bound residues of Nsp1-C-ter during the different replica runs of 200 ns. (E) Average hydrogen bonds throughout the different replica of the simulation runs of Nsp1 and drugs-bound complexes.

**Figure 4 figure supplement 1:**
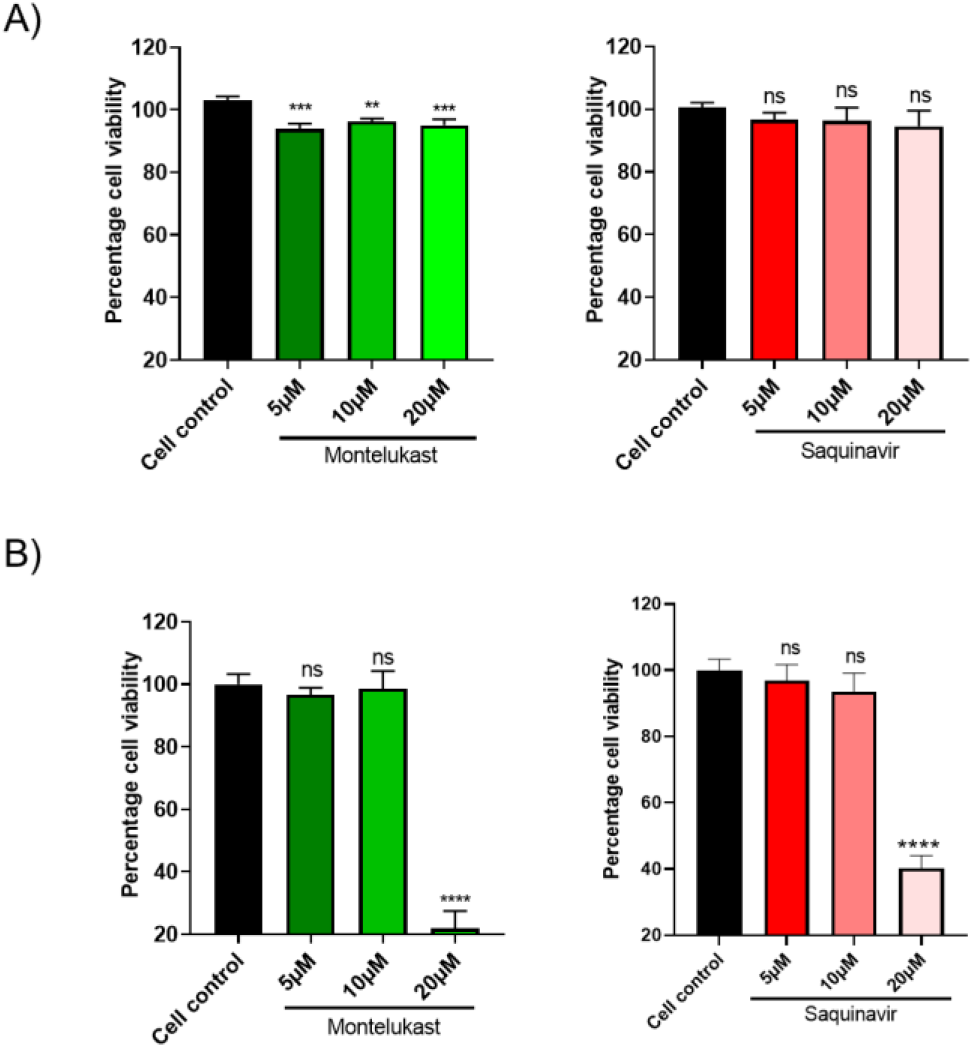
Cytotoxicity assay. Cells were treated in triplicates with increasing concentrations of montelukast or saquinavir as indicated, and cytotoxicity of the drugs was tested 48hr later by Alamar Blue assay. Data shows percentage toxicity of drugs compared to cell control in A) HEK-ACE2 and B) Vero E6 cells. **P < 0.01; ***P < 0.001; ****P < 0.0001; ns - not significant, using one-way ANOVA with Dunnett’s multiple comparison test. Error bars represent standard deviation.

**Figure 4 figure supplement 2.**
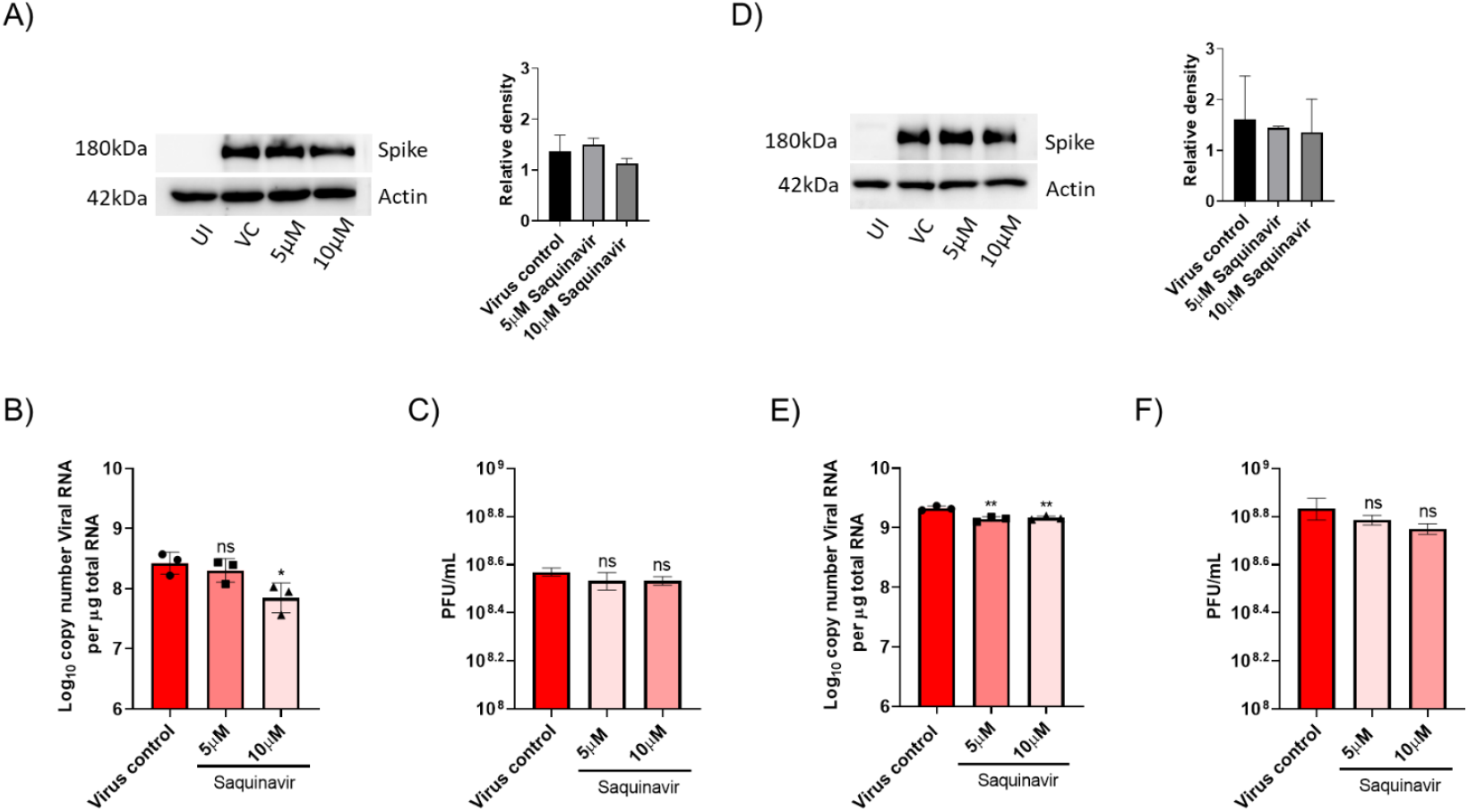
Saquinavir did not show significant antiviral activity against SARS-CoV-2. HEK ACE2 cells were pre-treated with 5 or 10 μM saquinavir and infected with 0.1 MOI SARS CoV-2 for 48hr. Virus spike protein expression by western blot analysis, with corresponding relative density of bands are shown in (A). Viral RNA copy number from infected cells was quantified by qRT PCR, and infectious virus titer from cell culture supernatants by plaque assay, shown in (B) and (C) respectively. Vero E6 cells were pre-treated with 5 or 10 μM saquinavir and infected with 0.001 MOI SARS CoV-2 for 48hr. (D) Virus spike protein expression by western blot analysis, with relative density of bands. (E) Viral RNA copy number from infected cells was quantified by qRT PCR and (F) infectious virus titer from cell culture supernatants by plaque assay. **P < 0.01; ***P < 0.001; ****P < 0.0001; ns - not significant, using one-way ANOVA with Dunnett’s multiple comparison test. Error bars represent standard deviation.

## References

Abu-Saleh, A.A.A., Awad, I.E., Yadav, A., and Poirier, R.A. (2020). Discovery of potent inhibitors for SARS-CoV-2’s main protease by ligand-based/structure-based virtual screening, MD simulations, and binding energy calculations. Physical chemistry chemical physics: PCCP 22, 23099–23106. http://www.ncbi.nlm.nih.gov/pubmed/33025993 PubMed Google Scholar

Babaylova, E., Malygin, A., Gopanenko, A., Graifer, D., and Karpova, G. (2019). Tetrapeptide 60-63 of human ribosomal protein uS3 is crucial for translation initiation. Biochimica et biophysica acta Gene regulatory mechanisms 1862, 194411. http://www.ncbi.nlm.nih.gov/pubmed/31356988 PubMed Google Scholar

Case, J.B., Bailey, A.L., Kim, A.S., Chen, R.E., and Diamond, M.S. (2020). Growth, detection, quantification, and inactivation of SARS-CoV-2. Virology 548, 39–48. http://www.ncbi.nlm.nih.gov/pubmed/32838945 PubMed Google Scholar

Cate, J.H. (2017). Human eIF3: from ‘blobology’ to biological insight. Philosophical transactions of the Royal Society of London Series B, Biological sciences 372. http://www.ncbi.nlm.nih.gov/pubmed/28138064 PubMed Google Scholar

Finkel, Y., Mizrahi, O., Nachshon, A., Weingarten-Gabbay, S., Morgenstern, D., Yahalom-Ronen, Y., Tamir, H., Achdout, H., Stein, D., Israeli, O., Beth-Din, A., Melamed, S., Weiss, S., Israely, T., Paran, N., Schwartz, M., and Stern-Ginossar, N. (2021). The coding capacity of SARS-CoV-2. Nature 589, 125–130. http://www.ncbi.nlm.nih.gov/pubmed/32906143 PubMed Google Scholar

Genheden, S., and Ryde, U. (2015). The MM/PBSA and MM/GBSA methods to estimate ligand-binding affinities. Expert opinion on drug discovery 10, 449–461. http://www.ncbi.nlm.nih.gov/pubmed/25835573 PubMed Google Scholar

Gordon, D.E., Jang, G.M., Bouhaddou, M., Xu, J., Obernier, K., White, K.M., O’Meara, M.J., Rezelj, V. V., Guo, J.Z., Swaney, D.L., Tummino, T.A., Huttenhain, R., Kaake, R.M., Richards, A.L., Tutuncuoglu, B., Foussard, H., Batra, J., Haas, K., Modak, M., Kim, M., Haas, P., Polacco, B.J., Braberg, H., Fabius, J.M., Eckhardt, M., Soucheray, M., Bennett, M.J., Cakir, M., McGregor, M.J., Li, Q., Meyer, B., Roesch, F., Vallet, T., Mac Kain, A., Miorin, L., Moreno, E., Naing, Z.Z.C., Zhou, Y., Peng, S., Shi, Y., Zhang, Z., Shen, W., Kirby, I.T., Melnyk, J.E., Chorba, J.S., Lou, K., Dai, S.A., Barrio-Hernandez, I., Memon, D., Hernandez-Armenta, C., Lyu, J., Mathy, C.J.P., Perica, T., Pilla, K.B., Ganesan, S.J., Saltzberg, D.J., Rakesh, R., Liu, X., Rosenthal, S.B., Calviello, L., Venkataramanan, S., Liboy-Lugo, J., Lin, Y., Huang, X.P., Liu, Y., Wankowicz, S.A., Bohn, M., Safari, M., Ugur, F.S., Koh, C., Savar, N.S., Tran, Q.D., Shengjuler, D., Fletcher, S.J., O’Neal, M.C., Cai, Y., Chang, J.C.J., Broadhurst, D.J., Klippsten, S., Sharp, P.P., Wenzell, N.A., Kuzuoglu-Ozturk, D., Wang, H.Y., Trenker, R., Young, J.M., Cavero, D.A., Hiatt, J., Roth, T.L., Rathore, U., Subramanian, A., Noack, J., Hubert, M., Stroud, R.M., Frankel, A.D., Rosenberg, O.S., Verba, K.A., Agard, D.A., Ott, M., Emerman, M., Jura, N., von Zastrow, M., Verdin, E., Ashworth, A., Schwartz, O., d’Enfert, C., Mukherjee, S., Jacobson, M., Malik, H.S., Fujimori, D.G., Ideker, T., Craik, C.S., Floor, S.N., Fraser, J.S., Gross, J.D., Sali, A., Roth, B.L., Ruggero, D., Taunton, J., Kortemme, T., Beltrao, P., Vignuzzi, M., Garcia-Sastre, A., Shokat, K.M., Shoichet, B.K., and Krogan, N.J. (2020). A SARS-CoV-2 protein interaction map reveals targets for drug repurposing. Nature 583, 459–468. http://www.ncbi.nlm.nih.gov/pubmed/32353859 PubMed Google Scholar

Jain, A.N. (2003). Surflex: fully automatic flexible molecular docking using a molecular similarity-based search engine. Journal of medicinal chemistry 46, 499–511. http://www.ncbi.nlm.nih.gov/pubmed/12570372 PubMed Google Scholar

Khan, A.R., Misdary, C., Yegya-Raman, N., Kim, S., Narayanan, N., Siddiqui, S., Salgame, P., Radbel, J., Groote, F., Michel, C., Mehnert, J., Hernandez, C., Braciale, T., Malhotra, J., Gentile, M.A., and Jabbour, S.K. (2021). Montelukast in hospitalized patients diagnosed with COVID-19. The Journal of asthma: official journal of the Association for the Care of Asthma, 1–7. http://www.ncbi.nlm.nih.gov/pubmed/33577360 PubMed Google Scholar

Ma, C., and Wang, J. (2021). Dipyridamole, chloroquine, montelukast sodium, candesartan, oxytetracycline, and atazanavir are not SARS-CoV-2 main protease inhibitors. Proceedings of the National Academy of Sciences of the United States of America 118. http://www.ncbi.nlm.nih.gov/pubmed/33568498 PubMed Google Scholar

Maier, J.A., Martinez, C., Kasavajhala, K., Wickstrom, L., Hauser, K.E., and Simmerling, C. (2015). ff14SB: Improving the Accuracy of Protein Side Chain and Backbone Parameters from ff99SB. Journal of chemical theory and computation 11, 3696–3713. http://www.ncbi.nlm.nih.gov/pubmed/26574453 PubMed Google Scholar

Min, Y.Q., Mo, Q., Wang, J., Deng, F., Wang, H., and Ning, Y.J. (2020). SARS-CoV-2 nsp1: Bioinformatics, Potential Structural and Functional Features, and Implications for Drug/Vaccine Designs. Frontiers in microbiology 11, 587317. http://www.ncbi.nlm.nih.gov/pubmed/33133055 PubMed Google Scholar

Narayanan, K., Huang, C., Lokugamage, K., Kamitani, W., Ikegami, T., Tseng, C.T., and Makino, S. (2008). Severe acute respiratory syndrome coronavirus nsp1 suppresses host gene expression,including that of type I interferon, in infected cells. Journal of virology 82, 4471–4479. http://www.ncbi.nlm.nih.gov/pubmed/18305050 PubMed Google Scholar

Paggiaro, P., and Bacci, E. (2011). Montelukast in asthma: a review of its efficacy and place in therapy. Therapeutic advances in chronic disease 2, 47–58. http://www.ncbi.nlm.nih.gov/pubmed/23251741 PubMed Google Scholar

Pettersen, E.F., Goddard, T.D., Huang, C.C., Couch, G.S., Greenblatt, D.M., Meng, E.C., and Ferrin, T.E. (2004). UCSF Chimera--a visualization system for exploratory research and analysis. Journal of computational chemistry 25, 1605–1612. http://www.ncbi.nlm.nih.gov/pubmed/15264254 PubMed Google Scholar

Sanghai, N., and Tranmer, G.K. (2020). Taming the cytokine storm: repurposing montelukast for the attenuation and prophylaxis of severe COVID-19 symptoms. Drug discovery today 25, 2076–2079. http://www.ncbi.nlm.nih.gov/pubmed/32949526 PubMed Google Scholar

Schmid, N., Eichenberger, A.P., Choutko, A., Riniker, S., Winger, M., Mark, A.E., and van Gunsteren, W.F. (2011). Definition and testing of the GROMOS force-field versions 54A7 and 54B7. European biophysics journal: EBJ 40, 843–856. http://www.ncbi.nlm.nih.gov/pubmed/21533652 PubMed Google Scholar

Schubert, K., Karousis, E.D., Jomaa, A., Scaiola, A., Echeverria, B., Gurzeler, L.A., Leibundgut, M., Thiel, V., Muhlemann, O., and Ban, N. (2020). SARS-CoV-2 Nsp1 binds the ribosomal mRNA channel to inhibit translation. Nature structural & molecular biology 27, 959–966. http://www.ncbi.nlm.nih.gov/pubmed/32908316 PubMed Google Scholar

Sharifulin, D.E., Bartuli, Y.S., Meschaninova, M.I., Ven’yaminova, A.G., Graifer, D.M., and Karpova, G.G. (2016). Exploring accessibility of structural elements of the mammalian 40S ribosomal mRNA entry channel at various steps of translation initiation. Biochimica et biophysica acta 1864, 1328–1338. http://www.ncbi.nlm.nih.gov/pubmed/27346718 PubMed Google Scholar

Sharma, T., Abohashrh, M., Baig, M.H., Dong, J.J., Alam, M.M., Ahmad, I., and Irfan, S. (2021). Screening of drug databank against WT and mutant main protease of SARS-CoV-2: Towards finding potential compound for repurposing against COVID-19. Saudi journal of biological sciences 28, 3152–3159. http://www.ncbi.nlm.nih.gov/pubmed/33649700 PubMed Google Scholar

Thoms, M., Buschauer, R., Ameismeier, M., Koepke, L., Denk, T., Hirschenberger, M., Kratzat, H., Hayn, M., Mackens-Kiani, T., Cheng, J., Straub, J.H., Sturzel, C.M., Frohlich, T., Berninghausen, O., Becker, T., Kirchhoff, F., Sparrer, K.M.J., and Beckmann, R. (2020). Structural basis for translational shutdown and immune evasion by the Nsp1 protein of SARS-CoV-2. Science 369, 1249–1255. http://www.ncbi.nlm.nih.gov/pubmed/32680882 PubMed Google Scholar

Tian, W., Chen, C., Lei, X., Zhao, J., and Liang, J. (2018). CASTp 3.0: computed atlas of surface topography of proteins. Nucleic acids research 46, W363–W367. http://www.ncbi.nlm.nih.gov/pubmed/29860391 PubMed Google Scholar

Tidu, A., Janvier, A., Schaeffer, L., Sosnowski, P., Kuhn, L., Hammann, P., Westhof, E., Eriani, G., and Martin, F. (2020). The viral protein NSP1 acts as a ribosome gatekeeper for shutting down host translation and fostering SARS-CoV-2 translation. Rna. http://www.ncbi.nlm.nih.gov/pubmed/33268501 PubMed Google Scholar

V’Kovski, P., Kratzel, A., Steiner, S., Stalder, H., and Thiel, V. (2021). Coronavirus biology and replication: implications for SARS-CoV-2. Nature reviews Microbiology 19, 155–170. http://www.ncbi.nlm.nih.gov/pubmed/33116300 PubMed Google Scholar

Vankadari, N., Jeyasankar, N.N., and Lopes, W.J. (2020). Structure of the SARS-CoV-2 Nsp1/5’-Untranslated Region Complex and Implications for Potential Therapeutic Targets, a Vaccine, and Virulence. The journal of physical chemistry letters 11, 9659–9668. http://www.ncbi.nlm.nih.gov/pubmed/33135884 PubMed Google Scholar

Wang, E., Sun, H., Wang, J., Wang, Z., Liu, H., Zhang, J.Z.H., and Hou, T. (2019). End-Point Binding Free Energy Calculation with MM/PBSA and MM/GBSA: Strategies and Applications in Drug Design. Chemical reviews 119, 9478–9508. http://www.ncbi.nlm.nih.gov/pubmed/31244000 PubMed Google Scholar

Yuan, S., Peng, L., Park, J.J., Hu, Y., Devarkar, S.C., Dong, M.B., Shen, Q., Wu, S., Chen, S., Lomakin, I.B., and Xiong, Y. (2020). Nonstructural Protein 1 of SARS-CoV-2 Is a Potent Pathogenicity Factor Redirecting Host Protein Synthesis Machinery toward Viral RNA. Molecular cell 80, 1055–1066 e1056. http://www.ncbi.nlm.nih.gov/pubmed/33188728 PubMed Google Scholar

